# Extrachromosomal DNA-mediated glutamine synthetase 2 (*GS2*) amplification enables glufosinate resistance in *Amaranthus palmeri*

**DOI:** 10.1101/2024.12.13.628355

**Authors:** Matheus M Noguera, Patrice S Albert, Christopher A Saski, James A Birchler, Aimone Porri, Ingo Meiners, Jens Lerchl, Nilda Roma-Burgos

## Abstract

**Background:** Palmer amaranth achieves resistance to glufosinate by overproducing the chloroplastic glutamine synthetase (GS2) protein, a result of the amplification and overexpression of its nuclear coding gene. This study examined how amplified *GS2* copies are inherited, identified their physical location in the cell, and investigated the mechanism of *GS2* amplification.

**Results:** Segregation analysis revealed that inheritance of amplified *GS2* copies deviates from classical Mendelian patterns, with poor correlation between plant level resistance and *GS2* amplification. Fluorescence *in situ* hybridization revealed chromosomal insertions of *GS2* and potential extrachromosomal circular DNA (eccDNA), and variability in *GS2* amplification both among individual plants and within cells (not all cells in plants with high *GS2* copy number showed GS2 amplification). The unpredictable inheritance patterns and distribution of *GS2* copies across multiple chromosomes suggest a role for eccDNA in *GS2* amplification. This was confirmed through eccDNA sequencing, which also identified multiple isoforms of *GS2*.

**Conclusion:** This is the second documented case of herbicide resistance conferred by eccDNA-mediated target-site gene amplification in Palmer amaranth.

## 1 INTRODUCTION

*Amaranthus palmeri* is a troublesome weed with worldwide distribution and can significantly reduce yield of row crops if not properly managed.^1^ One of the key traits that contribute to the exceptional weediness of this species is its high genetic diversity, which enhances its ability to evolve herbicide resistance. Resistance to at least nine herbicide modes of action has been reported, which ranks Palmer amaranth second highest in resistance problems, behind only *Lolium rigidum* and *Poa annua*, with resistance to 12 modes of action.^2^

The latest instance of herbicide resistance in Palmer amaranth involves evolved resistance to glufosinate-ammonium (GFA), the only commercially available herbicide that inhibits glutamine synthetase (GS), an enzyme involved in nitrogen metabolism. Currently in the USA, populations from Missouri, North Carolina, and Arkansas were confirmed to be resistant to GFA.^3–5^ In the Missouri population, resistance was attributed to the overproduction of GS2, the chloroplast-located isoform, by means of gene amplification and overexpression acting concomitantly, but not independently.^3^ In a population from Arkansas, *GS2* amplification was detected in four survivors of GFA applications, and *GS2* overexpression was detected in three nontreated plants from the same population.^6^ The full-length of the *GS2* gene was not obtained and sequencing of the partial gene fragment did not reveal mutations associated with resistance. Whether *GS2* amplification and overexpression happened in the same plant was not determined, nor was it determined if *GS2* overexpression leads to GS2 overproduction.^6^ The mechanism of resistance in GFA-resistant populations from North Carolina remains to be investigated.^5^

Herbicide resistance by target-site gene amplification is relatively infrequent. The first report dates to 2010, when 5-enolpyruvylshikimate-3-phosphate synthase (*EPSPS*) amplification in glyphosate-resistant Palmer amaranth was discovered by Gaines *et al*.^7^ The same mechanism was documented later in *L. perenne* ssp. *multiflorum*,^8^ *Bassia scoparia*,^9^ *A. spinosus*,^10^ *Eleusine indica*,^11^ *Bromus diandrus*,^12^ *A. tuberculatus*,^13^ *Chloris truncata*,^14^ *P. annua*,^15^ *Hordeum glaucum*,^16^ and *Salsola tragus.*^17^ A relatively less-known case, and the only one reported for this mode-of-action group, was amplification of Acetyl-CoA carboxylase (*ACCase*) in *Digitaria sanguinalis* cross-resistant to ACCase inhibitors.^18^

The physical location and genomic context of amplified gene copies in the plant genome can give hints about its heritability and mechanism of amplification.^19^ For example, *EPSPS* copies in *B. scoparia* are arranged in tandem at the telomeric regions of homologous chromosomes, which lead Jugulam *et al.* to suggest that amplification was due to unequal crossover.^9^ In addition, the close location of *EPSPS* copies resulted in single-locus inheritance.^9^ In *E. indica*, *EPSPS* copies are present in two pairs of homologous chromosomes, indicating a possible role of transposable elements in *EPSPS* amplification.^20^ In Palmer amaranth, *EPSPS* copies were found in several chromosomes,^7^ and amplification was later determined to be eccDNA-mediated.^21^ The apparent random distribution of *EPSPS* copies in Palmer amaranth genome has prevented scientists from fully understanding and predicting its inheritance.^22–24^

Due to its novelty, several aspects of GFA resistance in Palmer amaranth are still unresolved. This research aimed to elucidate two key aspects: the nature of amplification of *GS2* and the inheritance patterns of herbicide resistance.

## 2 MATERIALS AND METHODS

### 2.1 Plant materials and *GS2* copy number (CN) analysis

All experiments described herein were conducted with an *A. palmeri* population named MO20 #2 F1, hereafter called GFA-R, which was previously characterized regarding its response to GFA.^3^

Unless otherwise stated, seeds were sown in 50-cell trays filled with a commercial potting mix (ProMix LP15; Premier Tech Horticulture, Quakertown, PA, USA) and thinned to one plant per cell a week after planting. Plants were kept in a greenhouse maintained at 32/28 C day/night temperatures, with a 14-h photoperiod achieved by supplemental light. Herbicide applications were done using a benchtop sprayer equipped with 80 0067 flat fan nozzles calibrated to deliver 187 L ha^-1^ of spray mix at 275 kPa.

Leaf sample collection, DNA extraction and *GS2* CN assay were done as described elsewhere,^3^ the latter with slight modifications. To determine *GS2* CN, actin was used as the internal reference gene in duplex reactions and samples were run in duplicate. Each qPCR reaction was composed of 12.5 µL of 2x GoTaq qPCR Probe Master Mix (Promega Corporation, Madison, WI, USA), 2.5 µL of each primer (0.5 µM), 0.5 µL of each probe (0.2 µM), 100 ng of DNA and water to a final volume of 25 µL. Assays were conducted in a CFX96 Real-Time System (Bio-Rad Laboratories, Hercules, CA, USA) under the following thermoprofile: 95 C for 2 min followed by 40 cycles of 95 C for 15 s, and 60 C for 1 min. Fluorescence measurements were taken at the end of each amplification step, and the 2^-ΔCt^ was used to calculate relative *GS2* CN.^25^

### 2.2 Inheritance of amplified GS2 copies and GFA resistance

To study the inheritance pattern of amplified *GS2* copies, 35 nontreated plants from the GFA-R population were used. A single sample was collected from the youngest, fully expanded leaf of each plant. DNA was extracted and *GS2* CN was assessed. Selected plants were transplanted to 5-L pots and grouped according to parental pairs (Table 1). After maturity, female inflorescences were harvested, air dried and threshed, and seeds were cleaned with an air blower and stored in glass vials at room temperature.

**Table 1.** Crosses designed to determine the inheritance of *GS2* copies. CN was quantified via qPCR using a single tissue sample per plant, collected at the 10-leaf stage.

| Cross | *GS2:Actin* copy number |  |
| --- | --- | --- |
|  | ♀ | ♂ |
| RR-1 | 12 | 13 |
| RR-2 | 23 | 14 |
| SR | 3 | 13 |
| RS | 8 | 2 |
| SS-1 | 2 | 2 |
| SS-2 | 2 | 3 |

Two experiments were conducted with the seeds from designed crosses. In the first experiment, 50 plants (7- to 10-cm tall) from each cross were sprayed with a 1x rate of GFA (657 g ai ha^-1^, Liberty 280 SL, BASF Corporation, Research Triangle Park, NC, USA).

Ammonium sulfate was added at 10 g L^-1^. Living plants were counted at 15 days after application (DAA), and data were converted to survival percentage. The experiment was repeated, with 150 plants per cross. The second experiment aimed to quantify *GS2* CN in the offspring from each cross. A total of 250 nontreated plants were analyzed (Table 2). Tissue collection, DNA extraction and *GS2* CN quantification were done as previously described.

**Table 2.**
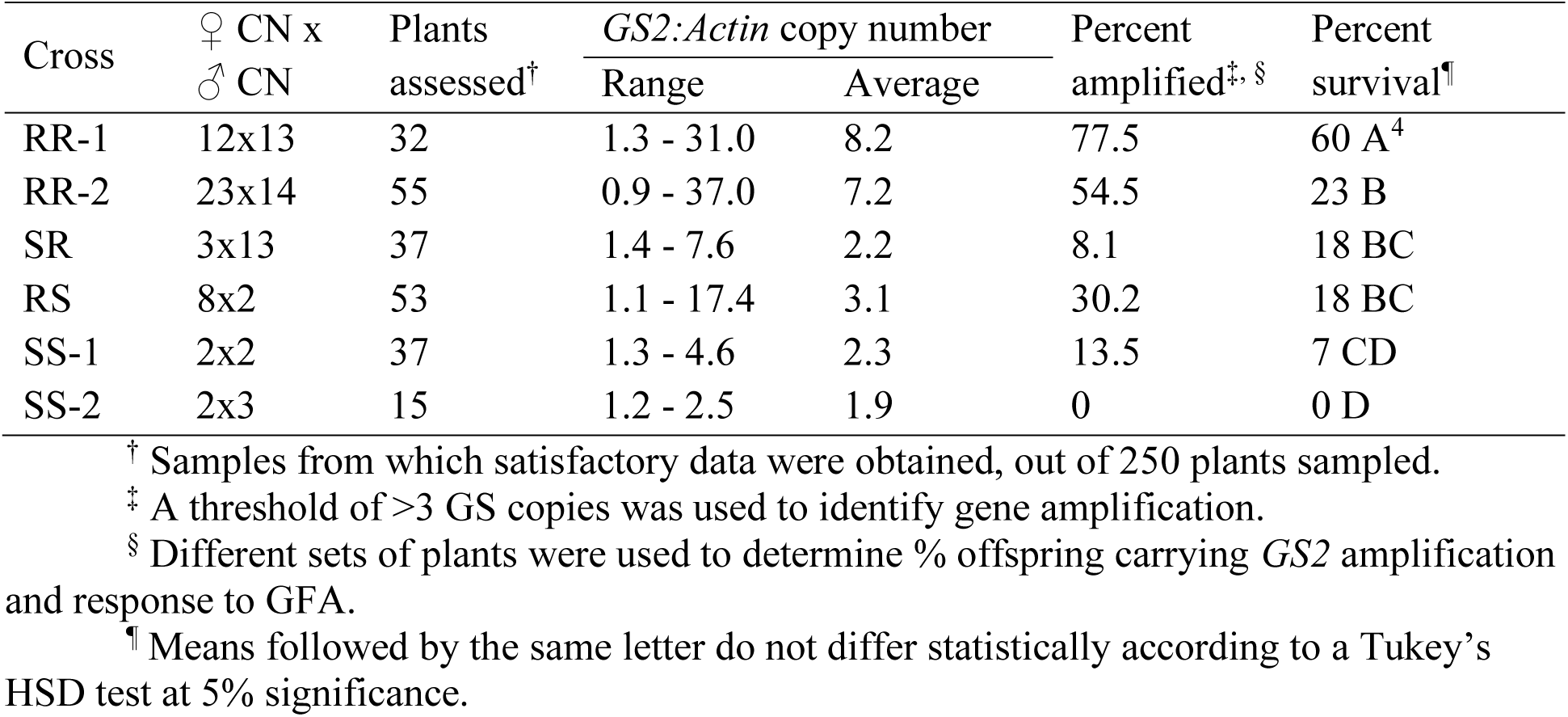
*GS2* CN in parent plants and offspring from six designed crosses, and response of offspring to GFA.

### 2.3 Fluorescence *in situ* hybridization (FISH)

Seeds from cross RR-1 (Table 1) were germinated in a tray containing potting mix, and 15 seedlings (1-leaf) were transferred into a hydroponic system. Plants were grown in a full-strength Hoagland basal salt solution, prepared by diluting a commercial salt mixture (MP Biomedicals LLC, Solon, OH, USA). The solution was aerated continuously using an aquarium air pump and replenished as needed. Three plants were selected for FISH assay, hereby designated as CN20, CN10 and CN1 based on their *GS2* CN, determined using DNA extracted from a single leaf per plant. Further examination of *GS2* CN on these plants was performed using DNA extracted from leaves and root samples.

To prepare the *GS2* FISH probe, a 2.8 kb region of the genomic sequence of *GS2* (full-length = 5.16 kb) was PCR-amplified using primers F: 5’-TGGCACAAATACTTGCACCTT-3’ and R: 5’-ACACTTGGGCCAACTTGGAA-3’. Genomic DNA from plant CN20 was used as a template for PCR amplification, with the following conditions: 98 C for 30 sec, followed by 30 cycles of 98 C for 10 sec, 60 C for 30 sec and 72 C for 3 min. A final extension step was added at the end of the run, being at 72 C for 5 min. The reaction consisted of 25 µL of EmeraldAmp Max HS Buffer, 1 µL of each primer (10 µM), 1 µL of DNA (50 ng µL^-1^) and 22 µL of nuclease-free water. Gel electrophoresis was performed to confirm amplicon size, and a commercial kit (PureLink Quick Gel Extraction Kit, ThermoFisher Scientific) was used to purify DNA fragments. Four identical PCR reactions were done, and purified DNA was bulked and concentrated to 200 ng µL^-1^. A *GS2* nick-translated probe was prepared using Texas Red-5- dCTP (PerkinElmer/Revvity, Boston, MA, USA) according to Kato *et al.*^26^ The telomeric probe, used to help delineate chromosomes, is a 5’ 6-FAM-labeled oligonucleotide, sequence 5’- TTTAGGGTTTAGGGTTTAGGGTTTAGGGTTTAGGG-3’ (Integrated DNA Technologies, Coralville, IA, USA).

Somatic chromosome spreads were prepared based on published protocols^27^ with minor modifications. Roots were digested in a mixture of 3% cellulase R-10 and 1.25% pectolyase Y23 (enzymes from Desert Biologicals Inc., Tempe, AZ, USA) for 15 to 20 min at 37 C after 1.5 h of N2O treatment at 1103 kPa. Chromosomes were counterstained with DAPI in Vectashield antifade solution (Vector Laboratories, Burlingame, CA, USA) and visualized with a BX61 fluorescence microscope (Olympus America Inc./Evident, Center Valley, PA, USA). Images were acquired with a mounted CCD camera using GenASIs software (Applied Spectral Imaging, Carlsbad, CA, USA) and processed using Adobe Photoshop 24.3.0 (Adobe Inc, San Jose, CA, USA).

### 2.4 Circular DNA enrichment and eccDNA sequencing

To determine if *GS2* copies are present in eccDNAs, two plants from the RR-2 population (Table 1) were selected with GS2 copy numbers of 24 and 37. Circular DNA was enriched and prepared for sequencing using the CIDER-Seq method by Mehta *et al*. ^28^ with a slight modification: the size exclusion step prior to enrichment was omitted. The enriched eccDNA from each sample was individually barcoded according to the manufacturer’s protocol (Pacific Biosciences, Menlo Park, CA, USA), pooled in equimolar amounts, and sequenced on a Sequel II single-molecule sequencer (Pacific Biosciences).

Raw sequence reads were demultiplexed and circular consensus sequences (CCS) were analyzed with the SMRT link software (Pacific Biosciences). Parameters for CCS analysis were stringent and included predicted quality = 0.999 and minimum read length = 1kbp. Processed reads were stored as fastq files, and those were analyzed with the packaged CIDER-seq software using the suggested approach to identify circular DNA. Predicted eccDNAs were matched to the *A. palmeri* reference genome.^29^ After processing of predicted eccDNA, shorter duplicate eccDNAs were collapsed into the longest reference eccDNA with the CDhit software,^30^ using an identity threshold of 90%. Reference eccDNA sequences were annotated for genuine open reading frames using the MAKER pipeline^31^ with evidence for genes derived from the *A. palmeri* published annotation.^29^ Alignments to the reference genome were performed with the Minimap2 software^32^ and comparative genome alignments performed with Mummer 4.0.^33^ Transfer RNAs were determined with the tRNAscan-SE software with default settings.^34^

## 3 RESULTS

### 3.1 Inheritance of amplified GS2 copies

To study the segregation patterns of amplified *GS2* copies, six crosses between individuals with known *GS2* CN were made. A total of 250 progenies were sampled (1 leaf sample per plant) and submitted to *GS2* CN analysis. Data were obtained for 229 plants. In addition, 200 seedlings per cross were sprayed with a labeled GFA rate and survival percentage was evaluated.

The progenies of RR-1 and RR-2 (described in Table 1) showed similar range and average of *GS2* CN. The highest *GS2* CN among siblings from these crosses was 31 and 37, respectively (Table 2). The average *GS2* CN was slightly higher in RR-1 siblings (8.2) than in RR-2 (7.2). The most remarkable difference between these two populations was the percentage of plants having *GS2* amplification: RR-1 had 23 percentage points higher compared to RR-2 (77.5 and 54.5%, respectively).

Crosses between individuals without *GS2* amplification are represented by SS-1 and SS-2. None of the 15 siblings analyzed from SS-2 showed more than 3 copies (Table 2). Conversely, 13% of the 37 siblings analyzed from SS-1 showed *GS2* amplification.

To determine possible gender-related effects in *GS2* segregation, populations SR and RS were created by crossing parents with and without *GS2* amplification. The RS cross had a high-copy female parent while the SR cross had a high-copy male parent. Interestingly, the RS progeny, which originated from a mother plant with *GS2* CN of 8, showed slightly higher average for this variable (3.1 copies) compared to SR progeny (2.2 copies). The percentage of RS siblings with *GS2* amplification was more than three times higher than that of the SR siblings, which in turn did not differ from SS-1, suggesting a possible maternal bias on inheritance of *GS2* amplification.

In terms of susceptibility to GFA, RR-1 and SS-2 crosses showed the highest and lowest survival percentages, respectively. The remaining crosses generated offspring that behaved similarly in response to GFA. Curiously, the survival percentage among progeny was lower than the percentage of individuals with *GS2* amplification in all cases but SR. In RR-2, for example, while 54.5% of seedlings showed *GS2* amplification, only 23% of the seedlings sprayed with GFA survived.

### 3.2 Physical mapping of amplified GS2 copies

To explore the physical location of *GS2* copies in Palmer amaranth plants, both with and without *GS2* amplification, fluorescence *in situ* hybridization (FISH) was conducted on the metaphase chromosomes and interphase nuclei of three plants exhibiting contrasting *GS2* CN. A plant telomeric DNA probe was included in the assay to enable quantitation of overlapping chromosomes and was detected in similar intensities in all 34 chromosomes.

In plants with *GS2* amplification, signals were observed in several mitotic chromosomes, with number and intensity of signals varying from cell to cell. For example, in plant CN20 signals were observed in up to ∼16 chromosomes (Figure 1A). Most of the *GS2* signals appeared in pairs, in a single-locus-gene fashion (as illustrated by Lamb *et al*.,^35^) although chromosomes containing 1 and >5 signals were also observed. Interestingly, only 11 out of 23 cells analyzed (or 48%) from plants CN20 and CN10 showed multiple *GS2* signals, while the remaining cells carried signals in a single chromosome pair (Figure 1 B).

**Figure 1.**
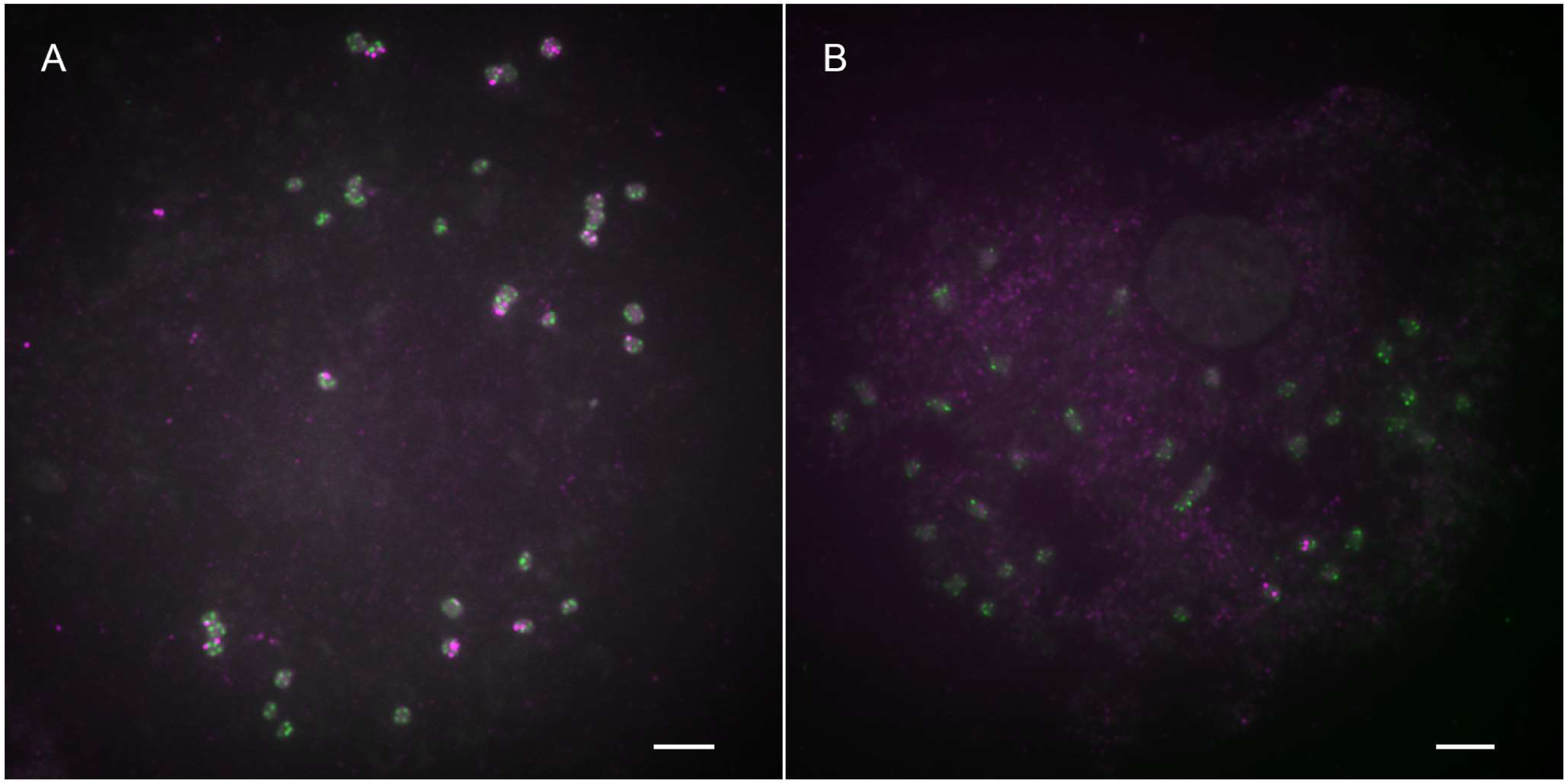
Variation in the number of *GS2*-FISH signals in a CN20 plant from a glufosinate-resistant population. (A) Root metaphase chromosome spread showing high *GS2* copy number. (B) Low GS2 copy number from a cell of the same plant. A telomere probe (green) was used to help define chromosome boundaries. Scale bars, 5 μm.

In plant CN1, originating from the same population as plants CN20 and CN10, but without *GS2* amplification detected by qPCR using DNA extracted from a leaf sample, 30 out of 32 cells displayed signals in a single chromosome pair (or two signals per interphase nuclei, as shown in Figure 2A). On the other hand, two spreads with signals in multiple chromosomes were observed (one shown in Figure 2B).

**Figure 2.**
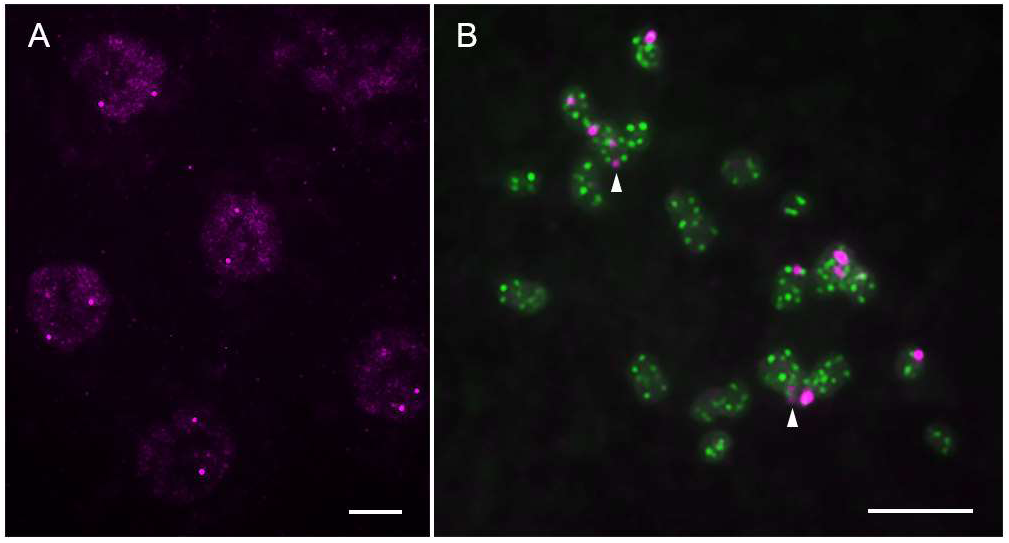
GS2 hybridization signals in a CN1 plant. **(A)** Nearly all cells had only a single copy of the gene (one pair of signals per nucleus). Hybridization channel is shown in magenta. **(B)** Root metaphase chromosomes from a cell showing numerous GS2 signals. Arrowheads denote hybridization to one pair of chromosomes. Telomere probe is shown in green. Scale bars, 5 μm.

After observing said results, further examination of *GS2* CN in these plants was performed using DNA extracted from leaf and root samples. The average *GS2* CN (n=4) in leaf and root tissues were, respectively: plant CN20 = 5.2 and 9.9; plant CN10 = 3.7 and 2.7; plant CN1 = 1.4 and 2.7 (Figure 3). When compared to our initial assessment, *GS2* amplification was present to a lower extent in plants CN20 and CN10. In plant CN1, virtually no variation was observed in *GS2* CN in DNA samples from leaves, but elevated number of copies was determined in two out of three root samples.

**Figure 3.**
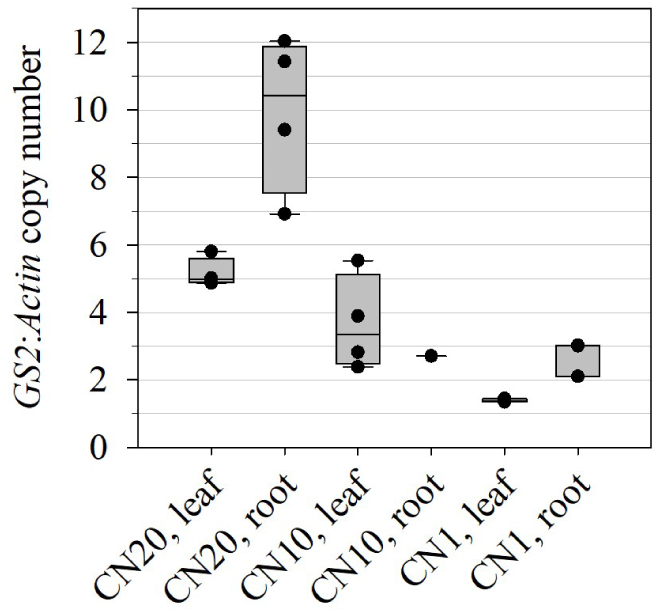
Box-whiskers plots of *GS2* CN in different tissues from plants used in the FISH experiment (n=4, with exception of CN10 root and CN1 root, where one and three samples were used, respectively). Dots represents individual data points.

Somatic mosaicism, the occurrence of genetically distinct lineages of cells within an organism derived from a single zygote, has been extensively researched, mainly in relation to human health.^36, 37^ As shown above, somatic mosaicism was present to varying degrees in the three plants studied, in both abundance of *GS2*-FISH signals and *GS2* CN. Similar results have been reported in glyphosate-resistant Palmer amaranth populations, using both genomic or cytogenetic approaches.^21, 24^ Thirty-nine percent of cells from plants with *GS2* amplification displayed multiple *GS2* signals, in contrast to only 6% of cells showing multiple signals in plant CN1, where *GS2* amplification was absent.

### 3.3 eccDNA sequencing

Using the CIDER-Seq methodology, we identified multiple isoforms of the *GS2* gene on eccDNAs of varying sizes. The *GS2* variants ranged from approximately 850 to 1300 bases in length and contained up to 13 exons. These isoforms were located on eccDNAs ranging from 5 to 21 kb in size (Table 3, Figure 4).

**Figure 4.**
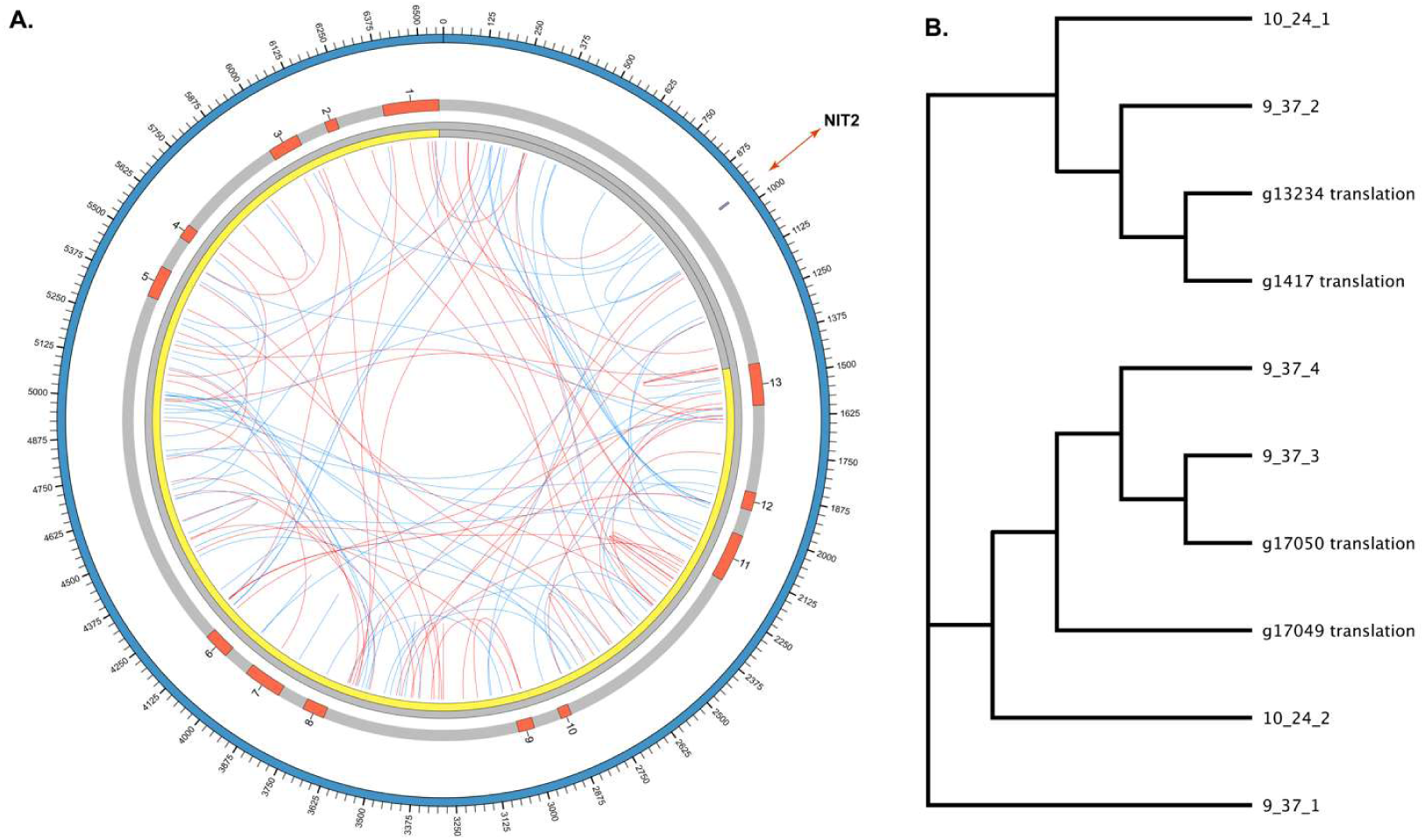
**(A)** Representative Circos plot of a *GS2*-encoding eccDNA. The outer blue ideogram represents the circular structure of the eccDNA with a double-headed arrow indicating the predicted NIT2 TF binding site. The next inner grey track shows the structure of the 13 predicted exons of the *GS2* focal amplification. The yellow track highlights the CDS regions. The red links show direct repeat matches while the blue links indicate inverted repeats of at least 10 bp and 90% identity. (**B)** A dendrogram of predicted *GS2* genes encoded on eccDNA and their relationship with published isoform sequences (*_translation).

**Table 3.**
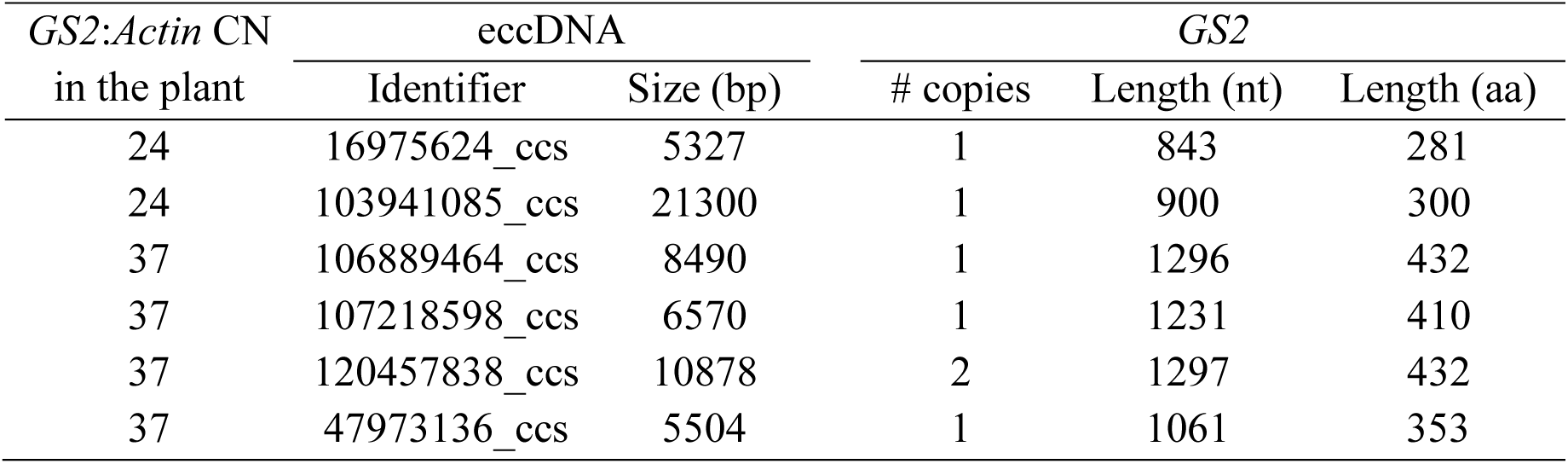
Summary of *GS2*-encoding eccDNAs.

Analysis of *GS2*-containing eccDNAs revealed a moderate level of repetitive complexity, indicating potential internal recombination sites. The eccDNA-encoded GS isoforms ranged in length from 281 to 431 amino acids (aa) and exhibited >95% sequence homology with *GS2* and *GS2.1* isoforms.^3^ Notably, a subset of the *GS* isoforms clustered with the cytosolic isoforms *GS1.1* and *GS1.2* of Palmer amaranth. This diversity among eccDNA-encoded GS sequences aligns with the high recombination rates and genetic heterogeneity characteristic of eccDNAs.

Sequence analysis of eccDNAs unveiled a NIT2 transcription factor binding sequence situated proximal to the *GS2* transcription start site, which is also found in the *GS2* native locus.

*NIT-2* is a key regulatory gene found in *Neurospora crassa*, especially active under nitrogen deficiency conditions.^38^ Functional orthologs have been described in other filamentous fungi such *Aspergillus nidulans*^39^ and *Fusarium graminearum*^40^, but its involvement in transcriptional regulation of *GS* expression in plants remains to be determined. Moreover, an eccDNA possessed a 131 aa predicted J-domain-containing protein (also known as DnaJ proteins) in addition to a *GS2* gene. DnaJ proteins compose a large family of heat shock proteins called Hsp40 and act mainly as regulators of Hsp70 proteins, which are associated with protein folding and assembly.^41^ Even though several Hsp40 proteins have been associated with stress tolerance in plants,^42, 43^ the functionality and role of this particular protein in GFA resistance is unclear and requires further studies.

## 4 DISCUSSION

Compared to other herbicide resistance mechanisms, such as target-site mutations and herbicide metabolism, target-site gene amplification is a rare occurrence. Understanding how gene amplification occurs and how these additional gene copies are transmitted to offspring is crucial for developing effective management and stewardship practices. The knowledge generated over the last 14 years since *EPSPS* amplification was first detected in *A. palmeri* provides a valuable framework for investigating new cases of herbicide resistance involving gene amplification.

Numerous researchers have investigated the segregation of glyphosate resistance and amplified *EPSPS* copies in Palmer amaranth, however, no consensus has been reached, and the segregation pattern remains unresolved. Gaines *et al*.,^7^ Mohseni-Moghadam *et al*. ^22^ and Chandi *et al*. ^23^ attributed resistance to a partially dominant trait under nuclear control, with only the latter observing a segregation pattern consistent with a single-gene inheritance model. Giacomini *et al*.^24^ observed *EPSPS* copies being inherited in a non-Mendelian pattern, with both positive and negative transgressive segregation, similar to findings by Gaines *et al*.^7^ This pattern was attributed to somatic mosaicism and clonal variation in *EPSPS* copy number. A maternal bias on *EPSPS* inheritance was also reported, resulting from facultative apomictic seed production.^44^

The pattern of *GS2* CN in our data resembles the *EPSPS* CN inheritance reported by Giacomini *et al*.^24^ but with added complexity, given that the proportion of offspring with *GS2* amplification and their survival frequency with GFA treatment can vary. For example, *GS2* amplification was observed in 54% of seedlings from cross RR-2, yet only 23% survived GFA treatment, a survival frequency not statistically distinct from crosses with one low-CN parent (SR and RS). This incongruence may be due to a physiological requirement for a minimum CN threshold to attain resistance. For instance, in glyphosate-resistant Italian ryegrass, all plants surviving a full dose of glyphosate contained at least 10 *EPSPS* copies and ∼15 *EPSPS* copies was the estimated threshold for survival in Palmer amaranth.^8, 45^ The relationship between *GS2* CN and the manifestation of resistance to GFA is further complicated by the co-existence of *GS2* overexpression, which does not correlate tightly with *GS2* CN. Lastly, the presence of additional resistance traits, such as the overexpression of stress-protection genes and herbicide-metabolism genes as described in *A. palmeri*^46^ and *E. indica*^47, 48^ cannot be ignored at this point. A follow-up experiment designed to confirm whether the apparent maternal bias on *GS2* inheritance holds true is needed. Thus far, the only published study on inheritance of GFA resistance in Palmer amaranth showed a similar pattern.^5^ Further research is necessary to clarify the mechanism of GFA resistance in Palmer amaranth and to assess the role of facultative apomixis in contributing to *GS2* segregation bias.

The unexpected segregation patterns of *GS2* can be attributed to significant cellular variability, as shown in the FISH assay. In high CN plants, we observed *GS2*-FISH signals of varying intensities across up to 16 chromosomes, although most cells (52%) displayed signals on only a single pair of homologous chromosomes. In the low CN plant, rare cells showed signals in multiple chromosomes. This aligns with previous data we generated, where tissue samples were collected over time from different leaves within a plant and a significant spatiotemporal variation in *GS2* CN was observed (Noguera 2024, unpublished). Likewise, plants used in the FISH assay showed variation in *GS2* CN across different tissues. Together, these observations suggest that the mechanism of *GS2* amplification in Palmer amaranth differs from the *EPSPS* amplification in *B. scoparia* and *A. tuberculatus.*^9, 13, 49^ Among those species, tandem amplification of *EPSPS* was found in only two chromosomes, with *A. tuberculatus* also presenting aneuploidy through an additional circular chromosome with multiple *EPSPS* copies. Aneuploidy does not appear to be involved in *GS2* amplification, as no cells with more than the expected 34 chromosomes^50^ were observed.

The observed somatic mosaicism in the presence and localization of *GS2*-FISH signals, along with the unpredictable inheritance patterns of amplified copies, suggests that eccDNAs are involved in *GS2* amplification. This behavior aligns with the findings of Koo *et al*.^21^ and Gaines *et al*.,^7^ who documented eccDNA-mediated *EPSPS* amplification in Palmer amaranth. The presence of multiple *GS2-*FISH signals in a low CN plant is also a common occurrence among species that develop eccDNA-mediated herbicide resistance, such as observed in glyphosate-resistant Italian ryegrass^51^ and Palmer amaranth.^21^ We confirmed this hypothesis by applying CIDER-Seq to high *GS2* CN plants, which revealed functional *GS2* variants in the circulome.

The eccDNA-encoded *GS2* sequences varied in length and composition, reflecting the dynamic nature of these nucleic acids. Interestingly, in most cells, *GS2* signals appeared as pairs, one in each sister chromatid, resembling a single-locus gene structure. This contrasts with Koo *et al.,*^21^ suggesting that some *GS2* copies have been integrated into the genome instead of relying on the chromosome-tethering mechanism for inheritance.

While the mechanisms of eccDNA integration into linear chromosomes in plants remain poorly understood,^52^ similar phenomena have been observed with *vasa* gene amplification in *Oreochromis niloticus,*^53^ *XylA* genes in *Saccharomyces cerevisiae,*^54^ and in several human oncogenes.^55^ This might explain the discrepancy between *GS2* copy number and expression levels reported in our previous study,^3^ as reintegration could occur in proximity to strong promoter motifs or genomic regions subject to epigenetic regulation. Consequently, *GS2* expression levels could vary significantly depending on the integration sites of the amplified copies.

It is important to highlight that the plants used in the FISH assay were grown under optimal conditions without exposure to GFA. In addition to the natural variability in eccDNA composition across tissues from a single organism,^56^ eccDNA abundance in plants was shown to vary in response to abiotic stress such as darkness or UV-light such as documented in rice.^57^ Therefore, it is possible that GFA application could affect both the proportion of cells with *GS2* amplification and the extent of amplification within those cells.^58^ Jugulam proposed that removing glyphosate selection pressure from resistant plants could eventually reduce *EPSPS* CN, potentially restoring susceptibility.^59^ However, this hypothesis has not been tested. In the case of *GS2* amplification, if our hypothesis that some eccDNAs integrate into linear chromosomes is correct, then removing glufosinate selection pressure would likely not restore population susceptibility.

The evolution of GFA resistance in Palmer amaranth through the complex mechanisms of *GS2* amplification and overexpression underscores the species’ remarkable adaptability. This finding further emphasizes the need for supplemental control methods to reduce reliance on herbicide use. Whether this mechanism of glufosinate resistance will evolve in other species, as seen with glyphosate resistance through *EPSPS* amplification,^60^ remains to be seen.

## 5 CONCLUSIONS

Inheritance of amplified *GS2* copies and GFA resistance deviates from Mendelian patterns. The correlation between the prevalence of *GS2* amplification and survival following GFA exposure is weak. *GS2* copies are distributed across multiple chromosomes and often appear integrated into the chromosome. The increase in GS2 copies is attributed to eccDNA, providing further evidence that these genomic elements play a prominent role in adaptive evolution.

## ACKNOWLEDGEMENTS

We thank James Heiser for providing the seeds of the original Palmer amaranth population used in this project and for being a collaborator and co-author of previous research on this population. This research was funded by BASF grant GR018328 to Nilda Roma-Burgos through the University of Arkansas Division of Agriculture, Agricultural Experiment Station.

## REFERENCES

1. Roberts J and Florentine S, A review of the biology, distribution patterns and management of the invasive species *Amaranthus palmeri* S. Watson (Palmer amaranth): Current and future management challenges. Weed Research 62: 113–122 (2022).

2. Heap I, International Survey of Herbicide Resistant Weeds. www.weedscience.org [accessed Jan 31 2024].

3. Noguera MM, Porri A, Werle IS, Heiser J, Brändle F, Lerchl J, et al., Involvement of glutamine synthetase 2 (GS2) amplification and overexpression in *Amaranthus palmeri* resistance to glufosinate. Planta 256: 1–14 (2022).

4. Priess GL, Norsworthy JK, Godara N, Mauromoustakos A, Butts TR, Roberts TL, et al., Confirmation of glufosinate-resistant Palmer amaranth and response to other herbicides. Weed Technology 36: 368–372 (2022).

5. Jones EAL, Dunne JC, Cahoon CW, Jennings KM, Leon RG and Everman WJ, Confirmation and inheritance of glufosinate resistance in an *Amaranthus palmeri* population from North Carolina. Plant-Environment Interactions 5: e10154 (2024).

6. Carvalho-Moore P, Norsworthy JK, González-Torralva F, Hwang J-I, Patel JD, Barber LT, et al., Unraveling the mechanism of resistance in a glufosinate-resistant Palmer amaranth (*Amaranthus palmeri*) accession. Weed Science 70: 370–379 (2022).

7. Gaines TA, Zhang W, Wang D, Bukun B, Chisholm ST, Shaner DL, et al., Gene amplification confers glyphosate resistance in *Amaranthus palmeri*. Proceedings of the National Academy of Sciences 107: 1029–1034 (2010).

8. Salas RA, Dayan FE, Pan Z, Watson SB, Dickson JW, Scott RC, et al., EPSPS gene amplification in glyphosate-resistant Italian ryegrass (*Lolium perenne* ssp. *multiflorum*) from Arkansas. Pest Management Science 68: 1223–1230 (2012).

9. Jugulam M, Niehues K, Godar AS, Koo D-H, Danilova T, Friebe B, et al., Tandem Amplification of a Chromosomal Segment Harboring 5-Enolpyruvylshikimate-3- Phosphate Synthase Locus Confers Glyphosate Resistance in *Kochia scoparia*. Plant Physiology 166: 1200–1207 (2014).

10. Nandula VK, Wright AA, Bond JA, Ray JD, Eubank TW and Molin WT, EPSPS amplification in glyphosate-resistant spiny amaranth (*Amaranthus spinosus*): a case of gene transfer via interspecific hybridization from glyphosate-resistant Palmer amaranth (*Amaranthus palmeri*). Pest Management Science 70: 1902–1909 (2014).

11. Chen J, Huang H, Zhang C, Wei S, Huang Z, Chen J, et al., Mutations and amplification of EPSPS gene confer resistance to glyphosate in goosegrass (*Eleusine indica*). Planta 242: 859–868 (2015).

12. Malone JM, Morran S, Shirley N, Boutsalis P and Preston C, EPSPS gene amplification in glyphosate-resistant Bromus diandrus. Pest Management Science 72: 81–88 (2016).

13. Dillon A, Varanasi VK, Danilova TV, Koo D-H, Nakka S, Peterson DE, et al., Physical mapping of amplified copies of the 5-enolpyruvylshikimate-3-phosphate synthase gene in glyphosate-resistant *Amaranthus tuberculatus*. Plant Physiology 173: 1226–1234 (2017).

14. Ngo TD, Malone JM, Boutsalis P, Gill G and Preston C, EPSPS gene amplification conferring resistance to glyphosate in windmill grass (*Chloris truncata*) in Australia. Pest Management Science 74: 1101–1108 (2018).

15. Brunharo CADCG, Morran S, Martin K, Moretti ML and Hanson BD, EPSPS duplication and mutation involved in glyphosate resistance in the allotetraploid weed species *Poa annua* L. Pest Management Science 75: 1663–1670 (2019).

16. Adu-Yeboah P, Malone JM, Gill G and Preston C, Non-Mendelian inheritance of gene amplification-based resistance to glyphosate in *Hordeum glaucum* (barley grass) from South Australia. Pest Management Science 77: 4298–4302 (2021).

17. Yanniccari M, Palma-Bautista C, Vázquez-García JG, Gigon R, Mallory-Smith CA and De Prado R, Constitutive overexpression of EPSPS by gene duplication is involved in glyphosate resistance in *Salsola tragus*. Pest Management Science 79: 1062–1068 (2023).

18. Laforest M, Soufiane B, Simard M-J, Obeid K, Page E and Nurse RE, Acetyl-CoA carboxylase overexpression in herbicide-resistant large crabgrass (*Digitaria sanguinalis*). Pest Management Science 73: 2227–2235 (2017).

19. Jugulam M and Gill BS, Molecular cytogenetics to characterize mechanisms of gene duplication in pesticide resistance. Pest Management Science 74: 22–29 (2018).

20. Chen J, Cui H, Ma X, Ma Y and Li X, Distribution differences in the EPSPS gene in chromosomes between glyphosate-resistant and glyphosate-susceptible goosegrass (*Eleusine indica*). Weed Science 68: 33–40 (2020).

21. Koo D-H, Molin WT, Saski CA, Jiang J, Putta K, Jugulam M, et al., Extrachromosomal circular DNA-based amplification and transmission of herbicide resistance in crop weed *Amaranthus palmeri*. Proceedings of the National Academy of Sciences 115: 3332–3337 (2018).

22. Mohseni-Moghadam M, Schroeder J and Ashigh J, Mechanism of Resistance and Inheritance in Glyphosate Resistant Palmer amaranth (*Amaranthus palmeri*) Populations from New Mexico, USA. Weed Science 61: 517–525 (2013).

23. Chandi A, Milla-Lewis SR, Giacomini D, Westra P, Preston C, Jordan DL, et al., Inheritance of evolved glyphosate resistance in a North Carolina Palmer amaranth (*Amaranthus palmeri*) biotype. International Journal of Agronomy 2012 2012).

24. Giacomini DA, Westra P and Ward SM, Variable inheritance of amplified EPSPS gene copies in glyphosate-resistant Palmer amaranth (*Amaranthus palmeri*). Weed Science 67: 176–182 (2019).

25. Schmittgen TD and Livak KJ, Analyzing real-time PCR data by the comparative CT method. Nature Protocols 3: 1101–1108 (2008).

26. Kato A, Kato A, Albert PS, Vega JM, Kato A, Albert PS, et al., Sensitive fluorescence in situ hybridization signal detection in maize using directly labeled probes produced by high concentration DNA polymerase nick translation. Biotechnic & Histochemistry 81: 71–78 (2006).

27. Kato A, Lamb JC and Birchler JA, Chromosome painting using repetitive DNA sequences as probes for somatic chromosome identification in maize. Proceedings of the National Academy of Sciences 101: 13554–13559 (2004).

28. Mehta D, Cornet L, Hirsch-Hoffmann M, Zaidi SS and Vanderschuren H, Full-length sequencing of circular DNA viruses and extrachromosomal circular DNA using CIDER-Seq. Nature Protocols 15: 1673–1689 (2020).

29. Montgomery JS, Giacomini D, Waithaka B, Lanz C, Murphy BP, Campe R, et al., Draft Genomes of *Amaranthus tuberculatus*, *Amaranthus hybridus*, and *Amaranthus palmeri*. Genome Biology and Evolution 12: 1988–1993 (2020).

30. Li W and Godzik A, Cd-hit: a fast program for clustering and comparing large sets of protein or nucleotide sequences. Bioinformatics 22: 1658–1659 (2006).

31. Cantarel BL, Korf I, Robb SM, Parra G, Ross E, Moore B, et al., MAKER: an easy-to- use annotation pipeline designed for emerging model organism genomes. Genome Research 18: 188–196 (2008).

32. Li H, Minimap2: pairwise alignment for nucleotide sequences. Bioinformatics 34: 3094–3100 (2018).

33. Marçais G, Delcher AL, Phillippy AM, Coston R, Salzberg SL and Zimin A, MUMmer4: A fast and versatile genome alignment system. PLOS Computational Biology 14: e1005944 (2018).

34. 34. Chan PP and Lowe TM. tRNAscan-SE: Searching for tRNA Genes in Genomic Sequences. In Gene Prediction: Methods and Protocols, ed. by Kollmar M. Springer New York: New York, NY, pp. 1-14 (2019).

35. Lamb JC, Danilova T, Bauer MJ, Meyer JM, Holland JJ, Jensen MD, et al., Single-gene detection and karyotyping using small-target fluorescence in situ hybridization on maize somatic chromosomes. Genetics 175: 1047–1058 (2007).

36. Forsberg LA, Gisselsson D and Dumanski JP, Mosaicism in health and disease — clones picking up speed. Nature Reviews Genetics 18: 128–142 (2017).

37. Youssoufian H and Pyeritz RE, Mechanisms and consequences of somatic mosaicism in humans. Nature Reviews Genetics 3: 748–758 (2002).

38. Fu Y-H and Marzluf GA, Characterization of nit-2, the major nitrogen regulatory gene of *Neurospora crassa*. Molecular and Cellular Biology 7: 1691–1696 (1987).

39. Davis MA and Hynes MJ, Complementation of areA-regulatory gene mutations of *Aspergillus nidulans* by the heterologous regulatory gene nit-2 of *Neurospora crassa*. Proceedings of the National Academy of Sciences 84: 3753–3757 (1987).

40. Giese H, Sondergaard TE and Sørensen JL, The AreA transcription factor in *Fusarium graminearum* regulates the use of some nonpreferred nitrogen sources and secondary metabolite production. Fungal Biology 117: 814–821 (2013).

41. Kampinga HH, Andreasson C, Barducci A, Cheetham ME, Cyr D, Emanuelsson C, et al., Function, evolution, and structure of J-domain proteins. Cell Stress and Chaperones 24: 7–15 (2019).

42. Cao L, Wang G, Fahim AM, Pang Y, Zhang Q, Zhang X, et al., Comprehensive Analysis of the DnaJ/HSP40 Gene Family in Maize (*Zea mays* L.) Reveals that ZmDnaJ96 Enhances Abiotic Stress Tolerance. Journal of Plant Growth Regulation 43: 1548-1569 (2024).

43. Liu J-Z and Whitham SA, Overexpression of a soybean nuclear localized type–III DnaJ domain-containing HSP40 reveals its roles in cell death and disease resistance. The Plant Journal 74: 110–121 (2013).

44. Ribeiro DN, Pan Z, Duke SO, Nandula VK, Baldwin BS, Shaw DR, et al., Involvement of facultative apomixis in inheritance of EPSPS gene amplification in glyphosate- resistant *Amaranthus palmeri*. Planta 239: 199–212 (2014).

45. Yakimowski SB, Teitel Z and Caruso CM, Defence by duplication: The relation between phenotypic glyphosate resistance and EPSPS gene copy number variation in *Amaranthus palmeri*. Molecular Ecology 30: 5328–5342 (2021).

46. Salas-Perez RA, Saski CA, Noorai RE, Srivastava SK, Lawton-Rauh AL, Nichols RL, et al., RNA-Seq transcriptome analysis of *Amaranthus palmeri* with differential tolerance to glufosinate herbicide. PLOS ONE 13: e0195488 (2018).

47. He S, Liu M, Chen W, Bai D, Liao Y, Bai L, et al., *Eleusine indica* Cytochrome P450 and Glutathione S-Transferase Are Linked to High-Level Resistance to Glufosinate. Journal of Agricultural and Food Chemistry 71: 14243–14250 (2023).

48. Lei T, Feng T, Wang L, Yuan X, Wu L, Wu B, et al., Metabolic resistance mechanism to glufosinate in *Eleusine indica*. Pesticide Biochemistry and Physiology 204: 106083 (2024).

49. Koo D-H, Jugulam M, Putta K, Cuvaca IB, Peterson DE, Currie RS, et al., Gene Duplication and Aneuploidy Trigger Rapid Evolution of Herbicide Resistance in Common Waterhemp Plant Physiology 176: 1932–1938 (2018).

50. Grant WF, Cytogenetic studies in *Amaranthus*.: III. Chromosome numbers and phylogenetic aspects. Canadian Journal of Genetics and Cytology 1: 313–328 (1959).

51. Koo D-H, Ju Y, Putta K, Sathishraj R, Roma-Burgos N, Jugulam M, et al., Extrachromosomal DNA-mediated glyphosate resistance in Italian ryegrass. Pest Management Science 79: 4290–4294 (2023).

52. Krasileva KV, The role of transposable elements and DNA damage repair mechanisms in gene duplications and gene fusions in plant genomes. Current Opinion in Plant Biology 48: 18–25 (2019).

53. Fujimura K, Conte MA and Kocher TD, Circular DNA Intermediate in the Duplication of Nile Tilapia vasa Genes. PLOS ONE 6: e29477 (2011).

54. Demeke MM, Foulquié-Moreno MR, Dumortier F and Thevelein JM, Rapid Evolution of Recombinant *Saccharomyces cerevisiae* for Xylose Fermentation through Formation of Extra-chromosomal Circular DNA. PLOS Genetics 11: e1005010 (2015).

55. Yang L, Jia R, Ge T, Ge S, Zhuang A, Chai P, et al., Extrachromosomal circular DNA: biogenesis, structure, functions and diseases. Signal Transduction and Targeted Therapy 7: 342 (2022).

56. Merkulov P, Egorova E and Kirov I, Composition and Structure of *Arabidopsis thaliana* Extrachromosomal Circular DNAs Revealed by Nanopore Sequencing. Plants 12: 2178 (2023).

57. Zhuang J, Zhang Y, Zhou C, Fan D, Huang T, Feng Q, et al., Dynamics of extrachromosomal circular DNA in rice. Nature Communications 15: 2413 (2024).

58. Arrey G, Keating ST and Regenberg B, A unifying model for extrachromosomal circular DNA load in eukaryotic cells. Seminars in Cell & Developmental Biology 128: 40–50 (2022).

59. Jugulam M, Can non-Mendelian inheritance of extrachromosomal circular DNA- mediated EPSPS gene amplification provide an opportunity to reverse resistance to glyphosate? Weed Research 61: 100–105 (2021).

60. Patterson EL, Pettinga DJ, Ravet K, Neve P and Gaines TA, Glyphosate Resistance and EPSPS Gene Duplication: Convergent Evolution in Multiple Plant Species. Journal of Heredity 109: 117–125 (2017).

